# A scalable Bayesian method for integrating functional information in genome-wide association studies

**DOI:** 10.1101/101691

**Authors:** Jingjing Yang, Lars G. Fritsche, Xiang Zhou, Gonçalo Abecasis, International Age-related Macular Degeneration Genomics Consortium (IAMDGC)

## Abstract

Although genome-wide association studies (GWASs) have identified many risk loci for complex traits and common diseases, most of the identified associations reside in noncoding regions and have unknown biological functions. Recent genomic sequencing studies have produced a rich resource of annotations that help characterize the function of genetic variants. Integrative analysis that incorporates these functional annotations into GWAS can help elucidate the biological mechanisms underlying the identified associations and help prioritize causal-variants. Here, we develop a novel, flexible Bayesian variable selection model with efficient computational techniques for such integrative analysis. Different from previous approaches, our method models the effect-size distribution and probability of causality for variants with different annotations and jointly models genome-wide variants to account for linkage disequilibrium (LD), thus prioritizing associations based on the quantification of the annotations and allowing for multiple causal-variants per locus. Our efficient computational algorithm dramatically improves both computational speed and posterior sampling convergence by taking advantage of the block-wise LD structures of human genomes. With simulations, we show that our method accurately quantifies the functional enrichment and performs more powerful for identifying true causal-variants than several competing methods. The power gain brought up by our method is especially apparent in cases when multiple causal-variants in LD reside in the same locus. We also apply our method for an in-depth GWAS of age-related macular degeneration with 33,976 individuals and 9,857,286 variants. We find the strongest enrichment for causality among non-synonymous variants (54x more likely to be causal, 1.4x larger effect-sizes) and variants in active promoter (7.8x more likely, 1.4x larger effect-sizes), as well as identify 5 potentially novel loci in addition to the 32 known AMD risk loci. In conclusion, our method is shown to efficiently integrate functional information in GWASs, helping identify causal variants and underlying biology.

**Author summary:** We propose a novel Bayesian hierarchical model to account for linkage disequilibrium (LD) and multiple functional annotations in GWAS, paired with an expectation-maximization Markov chain Monte Carlo (EM-MCMC) computational algorithm to jointly analyze genome-wide variants. Our method improves the MCMC convergence property to ensure accurate Bayesian inference of the quantifications of the functional enrichment pattern and fine-mapped association results. By applying our method to the real GWAS of age-related macular degeneration (AMD) with various functional annotations (i.e., gene-based, regulatory, and chromatin states), we find that the variants of non-synonymous, coding, and active promoter annotations have the highest causal probability and the largest effect-sizes. In addition, our method produces fine-mapped association results in the identified risk loci, two of which are shown as examples (*C2/CFB/SKIV2L* and *C3*) with justifications by haplotype analysis, model comparison, and conditional analysis. Therefore, we believe our integrative method will be useful for quantifying the enrichment pattern of functional annotations in GWAS, and then prioritizing associations with respect to the learned functional enrichment pattern.

## Introduction

Genome-wide association studies (GWASs) have identified thousands of genetic loci for complex traits and diseases, providing new insights into the underlying genetic architecture [1–5]. Each associated locus typically contains hundreds of variants in linkage disequilibrium (LD) [6,7], most of which are of unknown function and located outside protein-coding regions. Unsurprisingly, the biological mechanisms underlying the identified associations are often unclear [8] and pinpointing causal variants is difficult [9].

Recent functional genomic studies help understand and pinpoint causal variants and mechanisms [10–12]. Genetic variants can be annotated based on the genomic location (e.g., coding, intronic, and intergenic), role in determining protein structure and function (e.g., Sorting Intolerant From Tolerant (SIFT) [13] and Polymorphism Phenotyping (PolyPhen) [14] scores), ability to regulate gene expression (e.g., expression quantitative trait loci (eQTL) and allelic specific expression (ASE) evidence [15,16]), biochemical function (e.g., DNase I hypersensitive sites (DHS), metabolomic QTL (mQTL) evidence [17], and chromatin states [18–20]), evolutionary significance (e.g., Genomic Evolutionary Rate Profiling (GERP) annotations [21]), and a combination of different types of annotation (e.g., CADD [22]). Many statistical methods, including stratified LD score regression [23] and MINQUE [24], can now evaluate the role of functional annotations in GWASs through heritability analysis. Preliminary studies also show higher proportions of associated variants in protein-coding exons, regulatory regions, and cell-type-specific DHSs [25–27].

Integrating functional information into GWASs is expected to help identify and prioritize true causal associations. However, accomplishing this goal in practice requires methods to account for both LD and computational cost. Consider two recent methods, Fgwas [26] and PAINTOR [27], as examples: Fgwas assumes that variants are independent and there is at most one causal variant per locus, modeling no LD, which dramatically improves computational speed and allows Fgwas to be applied at genome-wide scale; PAINTOR accounts for LD, assuming the possibility of multiple association signals per locus, but is computationally slow and can only be used to fine-map small regions.

Here, we pair a flexible Bayesian method with an efficient computational algorithm. Together the two represent an attractive means to incorporate functional information into association mapping. Our model accounts for genotype correlation due to LD, allows for multiple causal variants per locus and, importantly, shares information genome-wide to increase association-mapping power. Our algorithm takes advantage of the local LD structure in the human genome [28–30] and refines previous Markov chain Monte Carlo (MCMC) algorithms to greatly improve mixing, which is key when searching for causal variants among many associated variants in LD (but less important in other applications such as modeling total genomic heritability). We refer to our method as the Bayesian functional GWAS (bfGWAS). Below, we illustrate the benefits of our method with extensive simulations and real data analyses of a large-scale GWAS on age-related macular degeneration (AMD) [31] with 33,976 individuals and 9,857,286 genotyped or imputed variants. The software for bfGWAS is freely available at https://github.com/yjingj/bfGWAS.

## Results

### Method overview

Our method is based on the standard Bayesian variable selection regression (BVSR) model (Online Methods and Text S1; Fig S1(a)), allowing for annotations that classify variants into *K* non-overlapping categories. We assume that variants in annotation category *q* share a “spike-and-slab” prior [32,33] for effect-sizes, 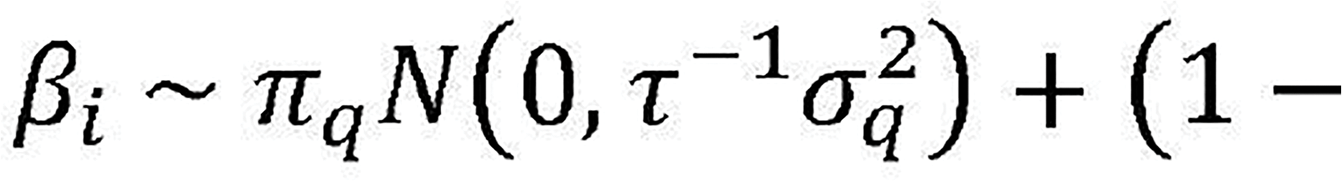

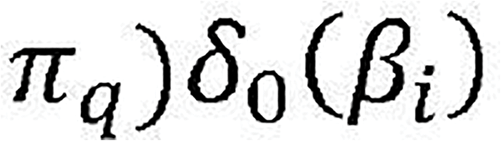. This model implies effect sizes are normally distributed as 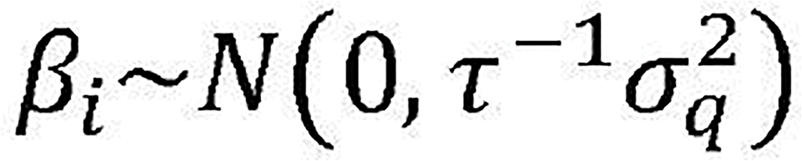 with probability *π*_*q*_, or set to zero with probability (1 - *π_q_*), with *δ*_0_(*β_i_*). denoting the point-mass function at 0. Here, *π*_*q*_represents the (unknown) causal probability for variants in the *q*th category and 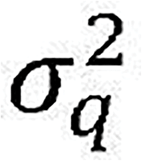 represents the (unknown) corresponding effect-size variance. An enhancement to previous Bayesian models [33–35] is that we model both the proportion of associated variants and their effect-size distribution in each annotation category.

Our goal is to simultaneously make inference on category specific parameters
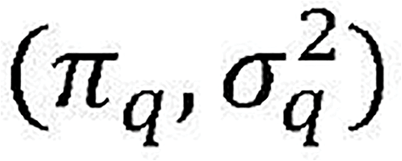 that represent the importance of each functional category, and on the variant specific parameters ––– effect-size *β*_*i*_ and the probability of *β*_*i*_ ≠ 0 (referred as posterior inclusion probability (*PP*_*i*_), representing association evidence). Our model shares information among genome-wide to estimate category specific parameters, which then inform the variant specific parameters. As a result, variant associations will be prioritized based on the inferred importance of functional categories.

Because standard MCMC algorithms suffer from heavy computational burden and poor mixing of posterior samples for large GWASs, we develop a novel scalable expectation-maximization MCMC (or EM-MCMC) algorithm. Our algorithm is based on the observation that LD decays exponentially with distance and displays local block-wise structure along the human genome [28–30,36,37]. This observation allows us to decompose the complex joint likelihood of our model into a product of block-wise likelihoods (Online Methods and Text S1). Intuitively, conditional on a common set of category specific parameters 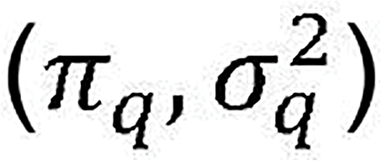, we can infer (*β*_*i*_, *PP*_*i*_), by running the MCMC algorithm per genome-block. A diagram of this EM-MCMC algorithm is shown in Fig S1(b).

Running MCMC per genome-block facilitates parallel computing and reduces the search space. Unlike previous MCMC algorithms for GWAS that use proposal distributions based only on marginal association evidence (such as implemented in GEMMA [38]), our MCMC algorithm uses a proposal distribution that favors variants near the “causal” variants being considered in each iteration, and prioritizes among these neighboring variants based on their conditional association evidence (see Text S1). Our strategy dramatically improves the MCMC mixing property, encouraging our method to explore different combinations of potentially causal variants in each locus (Fig S2). In addition, we implemented memory reduction techniques that reduce memory usage up to 97%, effectively reducing the required physical memory from 120 GB (usage by GEMMA [38]) to 3.6 GB for a GWAS with ~33K individuals and ~400K genotyped variants (Online Methods and Text S1).

In practice, we segment the whole genome into blocks of 5,000 ~ 10,000 variants, based on marginal association evidence, genomic distance, and LD. We always ensure variants in LD (R^2^ >0.1) with significant signals (P-values <5 × 10^−8^) are in the same block (Online Methods). We first initialize category specific parameters 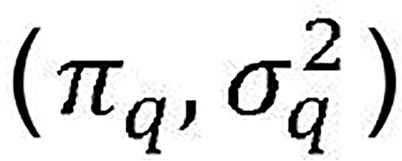, then run the MCMC algorithm per block (E-step), summarize the MCMC posterior estimates of (*β*_*i*_, *PP*_*i*_) across all blocks to update 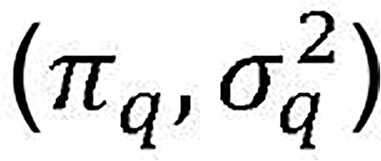(M-step), and repeat the block-wise EM-MCMC steps until 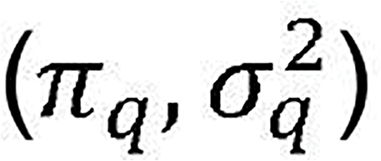 estimates converge (Fig S1(b)).

In addition, we calculate the regional posterior inclusion probability (regional-PP) per block that is the proportion of MCMC iterations with at least one “causal” variant (see Text S1). Because Bayesian PP might be split among multiple variants in high LD, the threshold of regional-PP >0.95 (conservatively analogous to false discovery rate 0.05) is used for identifying loci.

### Simulation

We simulated phenotypes with the genotype data (chromosomes 20-22) from the AMD GWAS [31], including 33,976 individuals and 241,500 variants with minor allele frequency (MAF) >0.1. We segmented this small genome into 50 × 2.5Mb blocks, each with ~5,000 variants. Within each block, we marked a 25KB continuous region (starting 37.5Kb from the beginning of a block) as the causal locus and randomly selected two causal single nucleotide polymorphisms (SNPs) per locus. We simulated two complementary annotations to classify variants into “coding” and “noncoding” groups, where the coding variants account for ~1% overall variants but ~10% variants within the causal loci (matching the pattern in the real AMD data). We simulated two scenarios: (i) coding variants ~44x enriched among causal variants (30 coding vs. 70 noncoding); (ii) no enrichment (1 coding vs. 99 noncoding). A total of 15% of phenotypic variance was divided equally among causal variants. We compared bfGWAS with single variant likelihood-ratio test, conditional analysis (CA), and Fgwas. The single variant test P-value (also referred to as P-value), conditioned P-value, Fgwas posterior association probability (PP, see Online Methods), and our Bayesian PP were used as criteria to identify associations.

We first compared power of different methods using average ROC curves[27,33] across 100 simulation replicates. Fgwas was more powerful than P-value at low false-positive rates (FPR), presumably because Fgwas incorporates annotation information (Fig 1(a)). However, with high false-positive rates, Fgwas underperformed P-value, presumably because Fgwas incorrectly assumes one variant per locus. In contrast, bfGWAS (modeling LD and allowing multiple causal variants per locus) outperformed both Fgwas and P-value for false-positive rates in (0, 0.01). Importantly, the advantage of bfGWAS became more pronounced with increasing sample size (Fig S3). Specifically, the power (based on FPR=0.5%) of bfGWAS increased from 48% to 64% as the sample size increased from 20K to 33K, while the power of Fgwas only increased from 52% to 56% and the power of P-values only increased from 47% to 52%. In addition, with sample size 33K and the threshold of regional-PP >0.95, bfGWAS has power 92.3% to identify associated loci, versus Fgwas with 88.6% power. The advantage of bfGWAS with large sample size suggests that bfGWAS can better extract the richer information available as sample size increases.

**Fig1.**
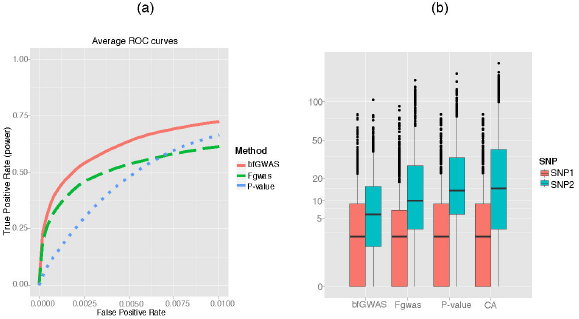
Power comparison among bfGWAS, Fgwas, P-value, and conditional analysis (CA), with 100 simulation replicates and the complete sample size 33,976. (a) Average ROC curves. (b) Boxplot of the ranks of the true causal SNP1 (with smaller P-value) and SNP2.

In a typical GWAS, researchers identify a series of associated loci and then examine associated variants within each locus independently. We examined the ability of each method to prioritize the true causal variants in each locus. Since we simulated two causal SNPs per locus (SNP1 and SNP2), we examine the power for identifying each of these separately (Fig 1(b)). All methods have the same median rank for causal SNP1 (typically, ranked 3rd rank among 150 SNPs in the locus by P-value, Fgwas and bfGWAS), suggesting that the strongest signal in a locus can often be identified without incorporating functional information. The median rank for the second causal SNP2 was the 7th by bfGWAS, 12th by Fgwas, 17th by P-value, and 18th by conditional analysis –– suggesting that incorporating functional information improves power to identify multiple signals in a locus. Stratified results based on the LD between two causal variants further demonstrate that bfGWAS has the highest power for identifying the weaker signal, especially when both SNPs are in high LD (Fig S4).

Both bfGWAS and Fgwas correctly identified enrichment in scenario (i) and properly controlled for the type I error of enrichment in scenario (ii), despite some numerical issues for Fgwas (Supplement Fig 5). Moreover, bfGWAS estimated the effect-size variance per annotation. For all 100 simulation replicates under both scenarios, the 95% confidence intervals of the log-ratio of estimated effect-size variances between coding and noncoding overlapped with 0 (Fig S6), suggesting effect-size variances were similar between two annotations (matching the simulated truth).

In summary, our simulation studies show that, in comparisons with competing methods, bfGWAS has higher power, especially in loci with multiple associated variants and when the sample size is large. Further, bfGWAS produces enrichment parameter estimates that can help with interpretation of association results.

### GWAS of AMD

Next, we applied our method to a GWAS of age-related macular degeneration (AMD) with 16,144 advanced cases and 17,832 controls, for a total of 33,976 unrelated European individuals. A total of 439,350 variants were genotyped on a customized Exome-Chip, and then imputed up to 12,023,830 variants in 1000 Genomes Project Phase 1 [39,40]. We analyzed 9,866,744 (~10M) low-frequency and common variants (MAF >0.5%) with three types of genomic annotations: gene-based functional annotations by SeattleSeq, summarized regulatory annotations [41], and the chromatin states profiled in nine human cell types from chromHMM [42,43].

### Coding variation and AMD

We used SeattleSeq to classify variants according to their impact on coding sequences (Table S1) and then applied our method bfGWAS and Fgwas. bfGWAS identified 37 loci out of 1,063 considered genome-blocks with regional-PP >0.95 (Tables S2, S3, and S5), including 32 among the 34 known AMD loci [31] and 5 potentially novel loci. Using the threshold of Bayesian PP >0.1068 (roughly equivalent to the P-value 5 × 10^−8^ based on permutations of AMD data; Fig S7), we identified 150 associated variants (Fig S9(a); Table S3), with 47 distributed among 42,005 non-synonymous variants, 4 among 67,165 synonymous coding variants, 54 among 3,679,235 intronic variants, 18 among 5,512,423 intergenic variants (including non-annotated variants), and 27 among 565,916 “other-genomic” variants (UTR, non-coding exons, upstream and downstream of genes). Very roughly, this corresponds to fraction of associated variants of ~1:1,000 among non-synonymous variants, 1:15,000 among synonymous variants, 1:100,000 among intronic variants, 1:300,000 among intergenic variants and 1:20,000 among “other-genomic” variants.

Similarly, Fgwas identified 46 loci by regional-PP >0.95, including all 34 known loci and 12 potentially novel loci (Tables S2, S4, and S6; Fig S9(b)). Since Fgwas analyzed the whole genome as 4,934 segments (each with 2,000 variants) and, thus, partitioned the genome somewhat differently than our method. Fgwas identified 178 associated variants with Fgwas PP >0.1068, including 24 non-synonymous, 13 coding-synonymous, 42 intronic, 40 intergenic, and 59 other-genomic signals. Compared with bfGWAS, the proportion of loci that contain at least one non-synonymous variant with PP >0.1068 is significantly smaller (11 out of 46 by Fgwas vs. 18 of 37 by bfGWAS; P-value = 0.017). Similarly, the proportion of non-synonymous variants prioritized by Fgwas is also significantly smaller (24 out of 178 by Fgwas vs. 47 of 150 by bfGWAS; P-value =7.7 × 10^−5^), indicating that bfGWAS places greater weight on coding variants ––– which, as a group, appears to have both a higher prior probability of association and larger effect sizes when associated.

Besides replicating the association results within known AMD loci[31], bfGWAS identified five novel loci (Table S5): missense *rs7562391/PPIL3*, *rs61751507/CPN1*, *rs2232613/LBP*, downstream *rs114348558/ZNRD1-AS1*, and splice *rs6496562/ABHD2*. These loci were also identified by Fgwas (Table S6) with different top association variants for *CPN1* (coding-synonymous *rs61733667*) and *ZNRD1-AS1* (downstream *rs116112857*). Interestingly, there are several connections between these potentially novel loci and known AMD loci. For example, the protein encoded by *LBP* is part of the lipid transfer protein family (which also includes *CETP* among the known AMD risk loci) that promotes the exchange of neutral lipids and phospholipids between plasma lipoproteins [44]. Similarly, *ZNRD1-AS1* has been associated with lipid metabolisms [45] and *ABHD2* has been associated with coronary artery disease [46], two other traits where the AMD loci encoding *CETP*, *APOE*, and *LIPC* are also involved. The gene *CPN1* has been associated with age-related disease (specifically, hearing impairment [47]).

**Multiple signals in a single locus**. We use two examples to illustrate the importance of studying multiple signals in a single locus. Our first example focuses on a 1Mb region around locus *C2/CFB/SKIV2L* on chromosome 6 where 1,862 variants have P-values < 5 ×10^−8^. There are an estimated 4 independent signals in the region by conditional analysis [31], 21 variants with Fgwas PP >0.1068, 11 with Bayesian PP >0.1068 by the standard Bayesian variable selection regression (BVSR) method that models no functional information, and 12 with Bayesian PP >0.1068 by bfGWAS. Interestingly, the alternative methods (P-value, Fgwas, and BVSR) identified intronic SNP *rs116503776/SKIV2L/NELFE* as the top candidates (P-value = 2.1 ×10^−114^; Fgwas PP = 0.912; BVSR PP = 1.0), while bfGWAS identified two missense SNPs *rs4151667/C2/CFB* (P-value = 1.4 × 10^−44^; bfGWAS PP = 0.917) and *rs115270436/SKIV2L/NELFE* (P-value = 2.8 × 10^−99^ SBA PP = 0.633) as the top functional candidates (Fig 2; Tables S2-S4).

**Fig2.**
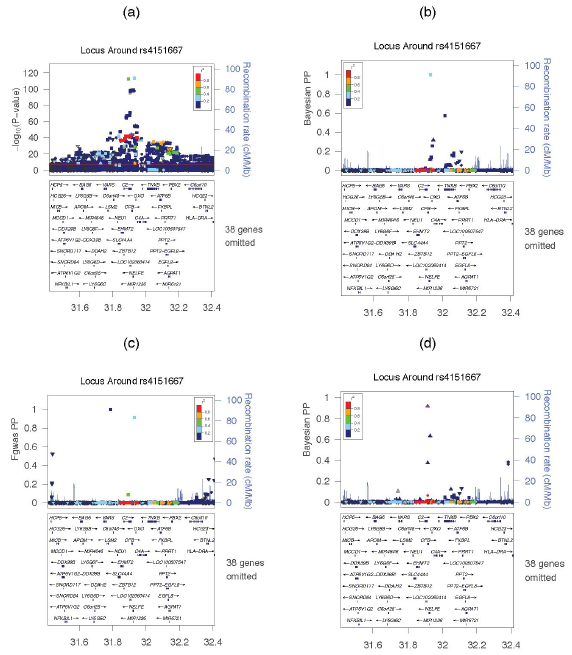
ZoomLocus plots around *rs4151667* in the locus *C2/CFB/SKIV2L* using the association criteria by P-value, standard BVSR, Fgwas, and bfGWAS. (a) −log10(P-values) by single variant tests. (b) Bayesian PPs by BVSR. (c) Fgwas PPs. (d) Bayesian PPs by bfGWAS. The top cyan squares in panels (a, b, c) denote the intronic variant *rs116503776*; the purple triangle in (d) denotes the non-synonymous variant *rs4151667*; shapes denote different annotations (triangle point up Δ for non-syn, circle ο for coding-syn, square ◻ for intronic, diamond ◊ for intergenic, and triangle point down ∇ for other-genomic).

A haplotype analysis describing the odds ratios (ORs) for all possible haplotypes for SNPs *rs116503776*, *rs4151667*, and *rs115270436*, helps clarify the region. Intronic SNP *rs116503776* with the smallest P-value appears to be associated with the phenotype by tagging the other two missense SNPs (Table S15). In particular, haplotypes with *rs116503776* can either increase or decrease risk, depending on alleles at the other two SNPs. To further confirm the importance of the missense SNPs *rs4151667* and *rs115270436*, we compared the AIC/BIC/loglikelihood between two models: one model with top two independent signals (*rs116503776* and *rs114254831*) identified by single-variant conditional analysis[31], versus the other model with top two signals (*rs4151667* and *rs115270436*) identified by bfGWAS. As expected, the second model has smaller AIC/BIC and larger loglikelihood than the first one (Table S16). Thus, we can see that while alternative methods (P-value, Fgwas, and BVSR) focus on the SNP with the smallest P-value, our bfGWAS method finds an alternative pairing of missense signals that better accounts for all data.

Our second example focuses on a 1Mb region around gene *C3* on chromosome 19 (Fig S10) with 112 genome-wide significant variants with P-value <5 ×10^−8^. Fgwas only discovered a single missense signal, *rs2230199* with the most significant P-value=1.7 ×10^−77^ (top blue triangle in Fig S10(a, c)). However, both BVSR and bfGWAS identified 2 missense variants with PPs = 1.0, and 5 intronic variants with 0.11< PPs <0.18. The top two missense signals *rs2230199* and *rs147859257* (241 base pairs apart) were confirmed by conditional analysis [31], where the second signal *rs147859257* has conditioned P-value=6.0 ×10^−33^ (the purple triangle in Fig S10(b, d), overlapping with *rs2230199*). These two missense signals match the interpretation of previous studies [48–50]. Because other 5 intronic variants (*rs11569479, rs11569470, rs201063729, rs10408682, rs11569466*) are in high LD with between variant R^2^ >0.98, we believe this is the third independent signal whose Bayesian PP was split among 5 variants in high LD by bfGWAS.

**Enrichment analysis.** bfGWAS estimated that non-synonymous variants are 10-100 times more likely to be causal than variants in other categories and that they also have larger effect-sizes (Fig 3(a, b)). To better compare enrichment among multiple categories, we define two new sets of parameters (Text S1). The first set of parameters, (*π*_*q*_/*π*_*avg*_), is defined to contrast the posterior association probability estimate (*π*_*q*_) for each category to the genome-wide average (*π*_*avg*_). The second set of parameters 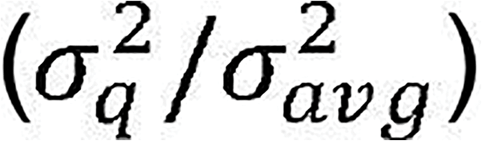 is similarly defined to contrast the effect-size variance from each category to the genome-wide average. Moreover, the square root of the effect-size variance reflects the effect-size magnitude because of the prior assumption for the effect-size in our model.

**Fig3.**
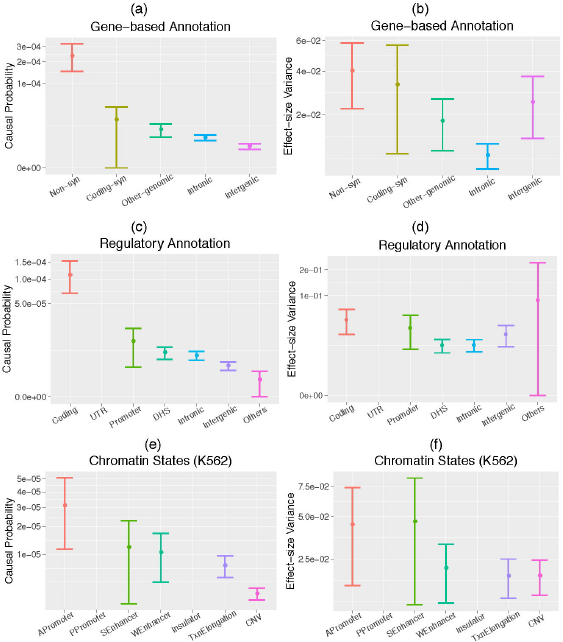
Category specific (enrichment) parameter estimates with 95% error bars by bfGWAS, with gene-based annotations, regulatory annotations, and chromatin-states profiled in the K562 cell line. (a, c, e) Causal probabilities. (b, d, f) Effect-size variances. The estimates that are the same as their priors are not plotted, e.g., estimates of UTR in (c, d), estimates of the active/poised promoter in (e, f). The estimate of the effect-size variance for the “Others” category in (d) is also close to the prior because of low region-association evidence, hence it has a wide 95% error bar.

Compared to the genome-wide average probability of causality *π*_*avg*_=4.3 ×10^−06^ (Fig S12(a)), we found that non-synonymous category were 54x more likely to be causal (P-value=7.24 ×10^−84^); that coding-synonymous and other variants were 4.3x and 2.2x more likely (P-values = 0.005, 0.003); and that intergenic 0.7x less likely (P-value=4.9 ×10^−6^); while the intronic variants matched the genome-wide average (P-value=0.659). In addition, compared to the genome-wide average effect-size variance 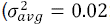; Fig S12(b)), we found that the effect size variance of was 1.9x larger for non-synonymous variants (P-value=0.014; i.e., 1.4x larger effect-size); and 0.4x smaller for variants in the intronic category (P-value=4.5 ×10^−06^); remaining categories were not significantly different (P-values >0.2). The estimated enrichment parameters by Fgwas show a similar pattern, although the contrast of the estimated enrichment for non-synonymous versus other annotations is not as pronounced as by bfGWAS (Fig S11(a)).

### Analysis with regulatory annotations

Second, we analyzed the GWAS data of AMD with the summarized regulatory annotations[41]: coding, UTR, promoter (defined as within 2KB of a transcription starting site), DHS in any of 217 cell types, intronic, intergenic, and “others” (not annotated as any of the previous six categories). Overall GWAS results were similar as the ones described in previous context (Tables S7-S10). Compared to the genome-wide average association probability (*π*_*avg*_=4.03 ×10^−6^ Fig S12(c)), we found that the association probability of the coding category was 28x higher (P-value < 2.2 ×10^−16^); the promoter was 2.6x (P-value=0.028) higher; the intergenic and “others” were 0.5x and 0.9x less (P-values = 5.3 ×10^−4^, 0.033); while the DHS and intronic were not significantly different (P-values>0.1). In addition, compared to the genome-wide average effect-size variance 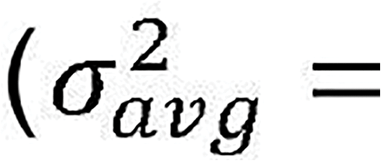 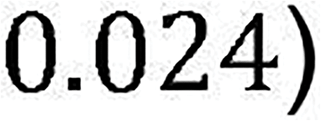, we found that the effect-size variance of the coding category was 1.9x larger (P-value=0.019; i.e., 1.4x larger effect-size); the DHS and intronic were 0.5x less (P-values = 0.011, 0.007); while the promoter, intergenic, and “others” were not significantly different (P-values >0.1; Fig S12(d)). Here, Fgwas identified a slightly different enrichment pattern (Fig S11(b)), where UTR was identified as the second most enriched category. This is presumably because Fgwas assumes one causal variant per locus and tends to prioritize the variant with the smallest P-value in each locus, e.g., UTR variants *rs1142/KMT2E/SPRK2* and *rs10422209/CNN2* have the highest Fgwas PP and the smallest P-value in their respective locus (Tables S2 and S8).

### Analysis with chromatin states

Last, we considered the annotations of seven chromatin states obtained with ChromHMM in nine human cell types[43]: active promoter (APromoter), poised promoter (PPromoter), strong enhancer (SEnhancer), weak enhancer (WEnhancer), insulator, transcription elongation (TxnElong), repetitive/copy number variation (CNV). Nine human cell types include: embryonic stem cells (H1-hESC), erythrocytic leukaemia cells (K562), B-lymphoblastoid cells (GM12878), hepatocellular carcinoma cells (HepG2), umbilical vein endothelial cells (HUVEC), skeletal muscle myoblasts (HSMM), normal lung fibroblasts (NHLF), normal epidermal keratinocytes (NHEK) and mammary epithelial cells (HMEC).

With each set of chromatin states profiled in one cell type, we applied bfGWAS on the GWAS data of AMD, and then examined the list of variants that contribute 95% posterior probabilities in the identified loci with regional-PP >95%. We found that the results by accounting for the chromatin states profiled in the erythrocytic leukaemia cells (K562) gave the shortest list (average 14 variants per locus; Table S17), and the enrichment analysis results of other cell types were slightly different (Figs S13–S15).

Here, we present the results of accounting for the chromatin states profiled in the K562 cell type (Fig 3(e, f); Tables S11-S14). Compared to the genome-wide average association probability (*π*_*avg*_4.0 ×10^−6^; Fig S12(e)), the association probability was 7.8x higher for the active promoter category (P-value = 7.4 ×10^−10^), 3x higher for the strong enhancer category (P-value=0.013), 2.6x higher for the weak enhancer category (P-value = 0.002), 1.8x higher for the transcription elongation category (P-value = 0.002), 0.4x less for the CNV category (P-values = 0.004). In addition, the effect-size variances of associated variants in active promoter and strong enhancer were found 2x larger than the genome-wide average 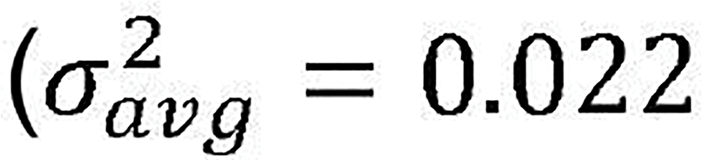; P-values = 0.048, 0.073), while the effect-size variances of weak enhancer, transcription elongation, and CNV categories were not significantly different (P-values >0.1; Fig S12(f)).

Note that the Bayesian enrichment estimates of the poised promoter and insulator categories are the same as their priors (not plotted in Fig 3(e, f)), suggesting that bfGWAS identified no associations in these two categories. Again, Fgwas identified a similar enrichment pattern (Fig S11(c)).

## Discussion

Here, we describe a scalable Bayesian hierarchical method, bfGWAS, for integrating functional information in GWASs to help prioritize functional associations and understand underlying genetic architecture. bfGWAS models both association probability and effect-size distribution as a function of annotation categories for improving fine-mapping resolution. Unlike previous methods [26,27], bfGWAS accounts for LD and allows for the possibility of multiple association signals per locus while remaining capable of genome-wide inference. Further, bfGWAS employs an improved MCMC sampling strategy to greatly improve the mixing of MCMC samples, which ensures the capability of identifying a list of association candidates.

By simulation studies, we demonstrated that bfGWAS had higher power than Fgwas and conditioned P-value for identifying multiple signals in a single locus by accounting for both functional information and LD. We also showed that bfGWAS accurately estimated the enrichment patterns under scenarios with or without enrichment for one annotation in simulations. In the real analysis using the AMD GWAS data and three different types of annotations, by bfGWAS, we obtained posterior association probabilities and effect-size variances for variants of considered annotation categories, as well as an improved list of fine-mapped association signals. In addition, we replicated the findings of 32 out of 34 known AMD risk loci, as well as identified 5 potentially novel loci by bfGWAS. Further, we gave two fine-mapped AMD loci *C2/CFB/SKIV2L* and *C3* by bfGWAS as examples with justifications by haplotype analysis, model comparison, and previous findings. Thus, we believe our method is useful for understanding the underlying genetic architecture of complex traits and diseases, for efficiently integrating functional information into GWASs.

Our flexible framework allows for many further extensions. For example, it can be extended to deal with overlapping or quantitative annotations (Text S1). These extensions will allow us to investigate the importance of a broader class of annotations (e.g. Combined Annotation Dependent Depletion (CADD) scores, MAF, and eQTL evidence). Importantly, as the development of new genomic assays and computational tools enables new variant annotations, simultaneous modeling of available annotations will be critical to identify the set of annotations that are important for a specific trait. Then extending bfGWAS to select relevant annotations would be useful.

bfGWAS makes a key assumption that the variant correlation matrix has a block-wise structure, which allows us to segment the genome into approximately independent blocks, analyze variants per block by MCMC, and summarize genome-wide information by an EM algorithm. In parallel to our study, many recent studies have also explored the benefits of dividing the human genome into approximately independent LD blocks to facilitate genome-wide analyses [26,51]. Although the standard segmentation methods (e.g., based on genomic location [51] as we adopted here, or the number of variants per block [26]) are often sufficient in practice, we expect that a better segmentation method [30] based on LD blocks will likely further increase the association mapping power.

The biggest limitation of bfGWAS is probably computational cost, as we perform MCMC using the complete genotype data. Specifically, bfGWAS took 5,000 CPU hours (~5 hours with parallel computations on 1,000 CPUs for the 1,063 genome-blocks) to analyze the AMD GWAS data with 33,976 individuals and 9,857,286 variants. Implementing bfGWAS with summary statistics is expected to reduce the computation cost significantly, which is part of our continuing project. In addition, the variational approximation [52,53] and other approximations [54,55] of MCMC may provide an efficient alternative for posterior inference in large GWAS.

## Materials and Methods

### Bayesian variable selection regression model

Our method is based on the standard Bayesian variable selection regression (BVSR) model

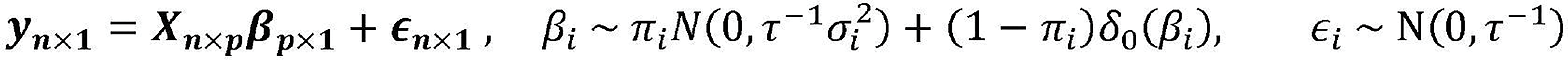,

where *n* denotes the number of individuals and *p* denotes the number of genetic variants; ***y*_*n×1*_** is the phenotype vector; ***X*_*n×p*_** is the genotype matrix; ***β*_*p*×1_** is a vector of genetic effect-sizes where each element *β_i_* follows a spike-and-slab prior (known as the point-normal distribution) ---- that is, *β*_*i*_ follows a normal distribution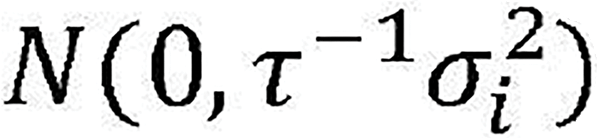 with probability *π*_*i*_, or *β*_*i*_ is set as 0 with probability (1 − *π*_*i*_) and a point mass density function *δ*_0_(*β*_*i*_) at 0 ( (*δ*_*0*_(*β*_*i*_)=1 if *β*_*i*_=0, *δ*_0_(*β*_*i*_)=0 otherwise) [32,33]; and *ε*_*i*_ is the residual error that independently and identically follows a normal distribution N(0, τ^-1^). We assume that both the phenotype vector ***y*_n×1_** and columns of the genotype matrix ***X*_*n*_*_×p_*** are centered, thus dropping the intercept. Although this model is developed for quantitative traits, we can treat binary phenotypes (e.g., cases and controls) as quantitative following previous approaches[33,35].

### Bayesian hierarchical model accounting for functional information

For integrating functional information into the above BVSR model, we classify all variants into disjoint categories by assuming one annotation per variant. We further assume that variants in the same functional category have the same spike-and-slab prior for the effect-sizes, i.e., 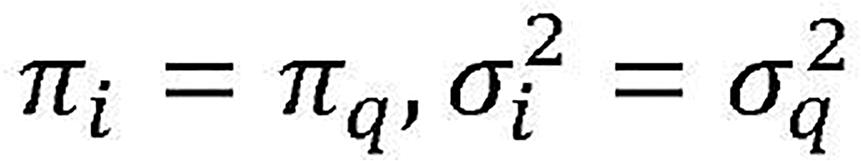 for the *q* th category. Consequently, *π_q_* denotes the category specific causal probability and 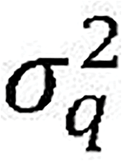 denotes the category specific effect-size variance (the square root of 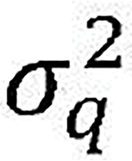 reflects the magnitude of effect size). Although we focus on discrete non-overlapping annotations in this paper, our method can be extended to overlapping and continuous annotations (Text S1).

We assume a Bayesian hierarchical framework[34] of BVSR with the following independent hyper priors:

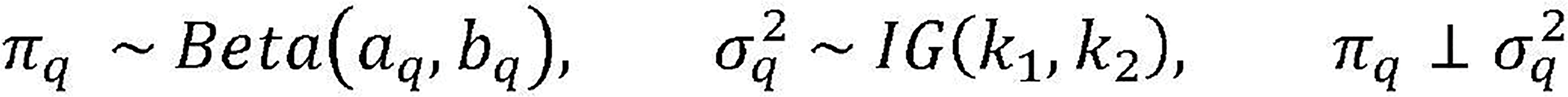

where *π*_*q*_ follows a Beta distribution with positive shape parameters *a*_*q*_ and *b*_*q*_, 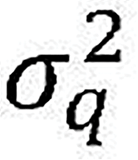 follows an Inverse-Gamma distribution with shape parameter *k*_1_ and scale parameter *k*_2_. In order to adjust for the unbalanced distribution of functional annotations among all variants and enforce a sparse model in our analysis, we choose values for *a*_*q*_ and *b*_*q*
_. such that the Beta distribution has mean 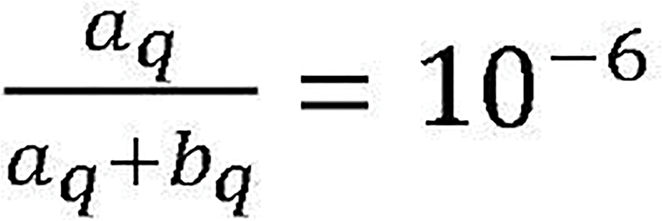 with 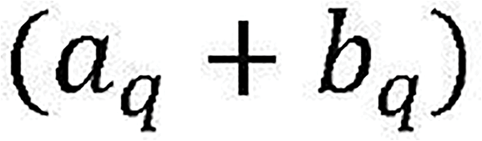 equal to the number of variants in category *q*. We set *k*_1_ =*k*_2_ =0.1 in our analysis to induce non-informative prior for 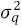. Note that τ is fixed at the phenotype variance value in our Bayesian inferences (Text S1).

### Bayesian references

We introduce a latent indicator vector ***γ*_*p×1*_** to facilitate computation, where each binary element *γ*_i_ indicates whether *β*_*i*_=0, by *γ*_*i*_=0, or 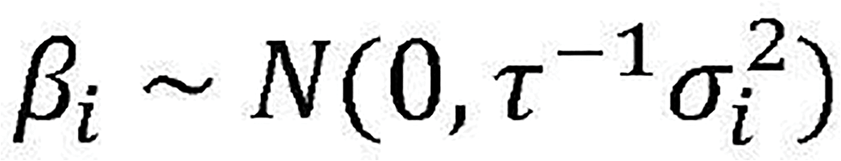 by *γ*_*i*_=1. Equivalently,

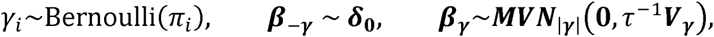

where **|*γ*|** denotes the number of 1’s in ***γ***; ***β*_−*γ*_** denotes the sub-vector of ***β*_*p×1*_** corresponding to variants with *γ*_*i*_;=0; ***β*_*i*_** denotes the sub-vector of ***β*_*p×1*_** corresponding to variants with (*γ*_*i*_;=0; j=1,., **|*γ*|**); and ***V*_*γ*_** denotes the sub-matrix of the diagonal matrix ***V*_*p×p*_** whose *ith* diagonal element is 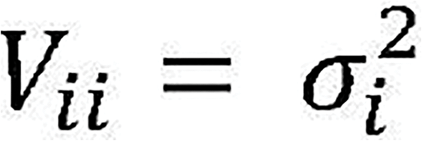. Consequently, the expectation of *γ*_*i*_is an estimate of the posterior inclusion probability (PP) for the *i*th variant, *E*[*γ_i_*]=*Prob(γ*_*i*_*=1)=PP*_i_.

For the described Bayesian hierarchical model above, the posterior joint distribution is proportional to

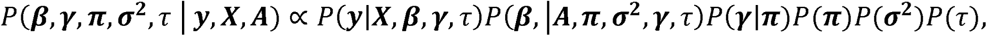

where 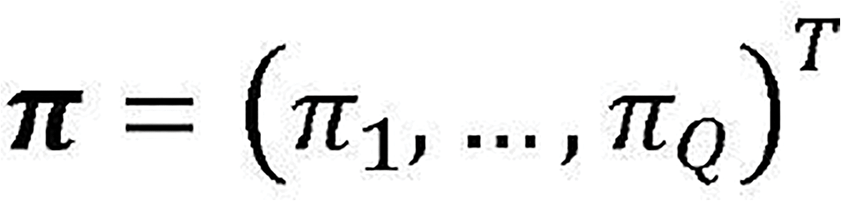, 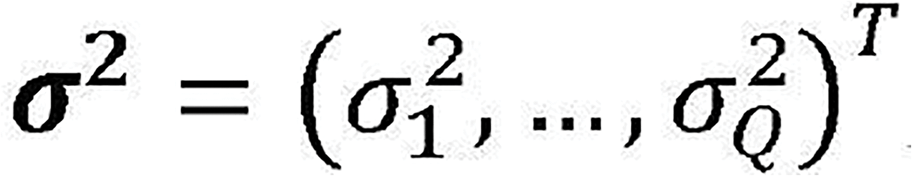, ***A*** is the *p × Q* matrix of binary annotations, and *Q* is the total number of annotations. The goal is to estimate the category specific parameters (***π*,*σ*^2^**) and the variant specific parameters (***β***, *E*[***γ***]) from their posterior distributions, conditioning on the data (***y, X, A***). Here, the category specific parameters denote the shared characteristics among all variants with the same annotation, which are also called enrichment parameters.

### EM-MCMC algorithm

The basic idea of the EM-MCMC algorithm is to segment the whole genome into approximately independent blocks each with 5,000 ~ 10,000 variants; run MCMC algorithm per block with fixed category specific parameter values (***π*,*σ*^2^**) to obtain posterior estimates of (***β***, *E*[***γ***]) (E-step); then summarize the genome-wide posterior estimates of (***β***, *E*[***γ***]) and update values of (***π*,*σ*^2^**) by maximizing their posterior likelihoods (M-step). Repeat such EM-MCMC iterations for a few times until the estimates of (***π*,*σ*^2^**) (maximum a posteriori estimates, i.e., MAPs) converge (Fig S1).

We derive the log-posterior-likelihood functions for (***π*,*σ*^2^**) and the analytical formulas for their MAPs. In addition, we construct their confidence intervals using Fisher information, whose analytical forms are derived for our Bayesian hierarchical model (Supplement Information). In our practical analyses, we find that, in general, with about 5 EM iterations, the estimates for (***π*,*σ*^2^**) would achieve convergence. Our method of conducting GWAS with functional information by using the above Bayesian hierarchical model and EM-MCMC algorithm is referred as “Scalable Functional Bayesian Association” (bfGWAS).

### Convergence diagnosis

Here, the MCMC algorithm is essentially a random walk over all possible linear regression models with combinations of variants, which can start with either a model containing multiple significant variants by sequential conditional analysis or the most significant variant by P-value. In each MCMC iteration, a new model is proposed by including an additional variant, or deleting one variant from the current model, or switching one variant within the current model with one outside; and then up to acceptation or rejection by the Metropolis-Hastings algorithm (Text S1). Importantly, we refine the standard proposal strategy for the switching step, by prioritizing variants in the neighborhood of the switch candidate according to their conditional association evidence (e.g., P-values conditioning on variants, except the switch candidate, in the current model). As a result, this MCMC algorithm encourages our method to explore different combinations of potentially causal variants in each locus, and significantly improves the mixing property.

We used the potential scale reduction factor (PSRF) [56] to quantitatively diagnose MCMC mixing property. PSRF is essentially a ratio between the average within-chain variance of the posterior samples and the overall-chain variance with multiple MCMC chains. From the example plots of the PSRFs of Bayesian PPs (Fig S2), for 58 top marginally significant SNPs (with P-values < 5 × 10^−8^) in the WTCCC GWAS data of Crohn’s disease [1], we can see that about half of the PSRF values by the standard MCMC algorithm (used in GEMMA [35]) exceed 1.2, suggesting the standard MCMC algorithm has poor mixing property. In contrast, the PSRF values by our MCMC algorithm are within the range of (0.9, 1.2), suggesting that our MCMC algorithm has greatly improved mixing property.

### Computational technics

We employ two computational technics to save memory in the bfGWAS software. One is to save all genotype data as unsigned characters in memory, because unsigned characters are equivalent to unsigned integers in (0, 256) that can be easily converted to genotype values within the range of (0.0, 2.0) by multiplying with 0.01. This technic saves up to 90% memory comparing with saving genotypes in double type. Second, with an option of in-memory compression, bfGWAS will further save additional 70% memory. As a result, we can decrease the memory usage from ~120 GB (usage by GEMMA[35]) to ~3.6GB, for a typical GWAS dataset with ~33K individuals and ~500K variants.

The bfGWAS software wraps a C++ executable file for the E-step (MCMC algorithm) and an R script for the M-step together by a Makefile, which is generated by a Perl script and enables parallel computation through submitting jobs. Generally, 50K MCMC iterations with ~5K variants and ~33K individuals take about 300MB memory and 1hr CPU time on a 1.6GHz core, where the computation cost is of order *O*(*nm*
^2^) with the sample size (*n*) and number of variants (*m*) considered in the linear models during MCMC iterations (usually *m* < 10). The computation cost for M-step is almost negligible because of analytical formulas for the MAPs.

### Fgwas

In this paper, the Fgwas results were generated by using summary statistics from single variant likelihood-ratio tests and the same annotation information used by bfGWAS. Fgwas[26] produces variant-specific posterior association probabilities (PPs), segment-specific PPs, and enrichment estimates for all annotations. To avoid the issue of failing convergence, we used segment size of 2,000 variants for Fgwas in both simulations and real data analyses. As a result, the final Fgwas PP is given by the product of the variant-specific PP and the corresponding segment–specific PP, and the Fgwas regional-PP is given by the highest segment-specific PP in a region or genome block.

**Simulation data**

We used genotype data on Chromosome 20-22 from the AMD GWAS (33,976 individuals and 241,500 variants with MAF>0.1) to simulate quantitative phenotypes from the standard linear regression model 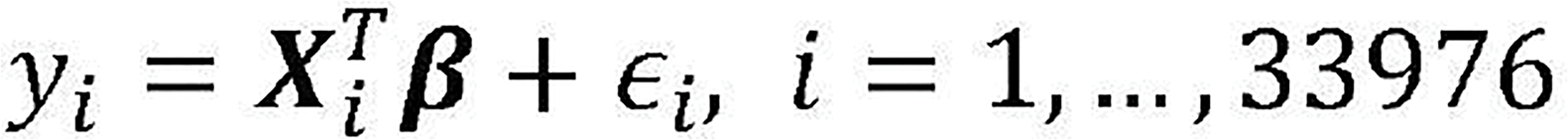, where ***X*_*i*_** is the genotype vector of the *ith* individual and ϵ*i* is the noise term generated from 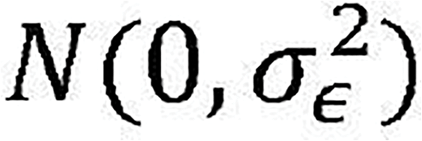. We segmented the genotype data into 50×2.5Mb blocks each with ~5,000 variants. Within each block, we marked a ~25Kb continuous region (starting 37.5Kb from the beginning of a block) as the causal locus and randomly selected two causal SNPs per locus. Two complementary annotations (“coding” vs. “noncoding”) were simulated, where the coding variants account for ~1% overall variants but ~10% variants within the causal loci (matching the pattern in the real AMD analysis). We selected positive effect-size vector ***β*** and noise variance 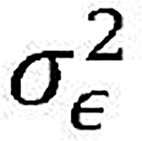 such that a total of 15% phenotypic variance was equally explained by causal SNPs. We controlled the enrichment-fold of coding variants by varying the number of coding variants among these 100 causal SNPs.

We compared bfGWAS with P-value, conditioned P-value, and Fgwas. In the simulation studies, P-values were obtained from a series of likelihood-ratio tests based on the standard linear regression model. P-values conditioning on the top significant variant per locus were used to identify the second signal by conditional analysis. Fgwas was implemented with summary statistics from single variant tests and the segment size of 2,000 variants (selected to avoid convergence issues). We failed to include PAINTOR in the comparison, because PAINTOR cannot complete the analysis for one block in >1,000 CPU hours (on a 2.5GHz, 64-bit CPU) and is thus expected to require >1 million CPU hours for a genome-wide analysis.

### GWAS data of AMD

In the GWAS data of AMD, the advanced AMD cases – including wet cases with choroidal neovascularization (CNV, when accompanied by angiogenesis) and dry cases with geographic atrophy (GA, when angiogenesis is absent) – and control subjects were gathered across 26 studies, with DNA samples collected and genotyped centrally [39]. All genotypes were generated by a customized chip that contains (i) the usual genome-wide variant content, (ii) exome content comparable to the Exome chip (protein-altering variants across all exons), (iii) variants in known AMD risk loci (protein-altering variants and previously associated variants), and (iv) previously observed and predicted variation in *TIMP3* and *ABCA4* (two genes implicated in monogenic retinal dystrophies). The genotyped variants (439,350) were then imputed to the 1000 Genomes reference panel (Phase I) [40], resulting a total of 12,023,830 variants.

The software bfGWAS used dosage genotype data and standardized phenotypes. Phenotypes were first coded quantitatively with 1’s for cases and 0’s for controls; second corrected for the first and second principle components, age, gender, and source of DNA samples; and then standardized to have mean 0 and standard deviation 1. In order to make the Bayesian inferences scalable to the AMD GWAS data (33,976 individuals, 9,866,744 variants with MAF >0.5%), we segmented the whole genome into 1,063 non-overlapped blocks, such that each block has length ~2.5Mb (containing ~10,000 variants) and all previously identified loci along with variants in LD (R^2^ >0.1) were not split. Then we applied the EM-MCMC algorithm with 5 EM steps and 50,000 MCMC iterations (including 50,000 extra burn-ins).

For comparison, P-values were obtained by a series of likelihood-ratio tests, using the same “quantitative” phenotype vector as used by bfGWAS; Fgwas was implemented with the summary statistics from single variant tests and the segment size of 2,000 variants (resulting 4,934 segments); and a standard Bayesian variable selection regression (BVSR) method that models no functional information was also applied.

Three types of genomic annotations were considered for analyzing the AMD data: gene-based functional annotations of SNPs and small indels from SeattleSeq (http://snp.gs.washington.edu/SeattleSeqAnnotation138/index.jsp), summarized regulatory annotations [41], and the chromatin states profiled respectively in nine human cell types from chromHMM [19,42,43]. For variants annotated with multiple functions, we used the most severe function in the analysis: non-synonymous > coding-synonymous > other-genomic > intronic > intergenic for the gene-based annotations; coding > UTR > promoter > DHS > intronic > intergenic > “others” for the summarized regulatory annotations; active promoter > poised promoter > strong enhancer > weak enhancer > insulator > transcription elongation > CNV for the chromatin states.

### Software

Our software bfGWAS is freely available on Github (https://github.com/yjingj/bfGWAS).

## Supporting information

Tables S1-S17.

Text S1. Supplementary text containing more details about the bfGWAS method.

Fig S1. Flowchart of the bfGWAS. (a) Bayesian hierarchical model. (b) Diagram of the EM-MCMC algorithm.

Fig S2. Plots of the PSRF values for the Bayesian PPs of 58 top marginally significant SNPs (using the WTCCC GWAS data of Crohn’s disease) with 3, 8, 15, and 20 MCMC chains. (a) Standard MCMC algorithm as in GEMMA. (b) Our MCMC algorithm in bfGWAS. PSRF within (0.9, 1.2) suggests good mixing property.

Fig S3. Average ROC curves of 100 simulation studies by P-value (single variant test), Fgwas, and bfGWAS with various sample sizes.

Fig S4. Prioritized ranks of the true causal SNP1 (pink) and SNP2 (cyan) by P-value (single variant test), conditional analysis (CA), Fgwas, and bfGWAS, stratified by the LD between SNP1 and SNP2. Our bfGWAS method outperforms the other three methods for identifying SNP2 in all scenarios, while the relative performance of the conditional analysis and Fgwas for identifying SNP2 depends on LD. In particular, when SNP1 and SNP2 are in low LD, the conditional analysis outperforms Fgwas. This presumably is due to the wrong Fgwas assumption of one causal per locus, and the reduced association strength for SNP2 by conditioning on the top significant variant (SNP1 or a proxy for SNP1) that is in high LD with SNP2.

Fig S5. Estimates of Fgwas enrichment parameters and log-relative-risk ln(*π*_0_/*π*_1_) by bfGWAS, along with 95% confidence intervals. (a) Simulation Scenario I with ~44x enrichment in coding. (b) Scenario II with no enrichment. No enrichment is estimated when the 95% confidence interval covers 0, while enrichment for coding is estimated with the 95% confidence interval above 0.

Fig S6. Estimates of the log-ratio of effect-size variances ln 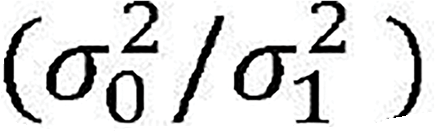 by bfGWAS, along with 95% confidence intervals. (a) Scenario I with ~44x enrichment in coding. (b) Scenario II with no enrichment. The effect-sizes of both groups in Scenarios I and II were simulated from the same normal distribution, thus the 95% confidence intervals covering 0 suggest that bfGWAS estimates similar effect-size variances between two categories.

Fig S7. Sorted top Bayesian PP by bfGWAS versus the sorted top -log10(P-values) of single variant tests for 100 GWASs with AMD genotype data and permuted phenotypes. Here, the P-value 5 × 10^−8^ roughly corresponds to Bayesian PP 0.1068.

Fig S8. Manhattan plot with -log10(P-values) by single variant tests for the GWAS of AMD, where variants with Bayesian PP >0.1068 by standard Bayesian variable selection regression (BVSR, modeling no functional information) are colored.

Fig S9. Manhattan plots with -log10(P-values) by single variant tests for the GWAS of AMD with gene-based annotations, where different shapes denote different functional annotations. (a) Variants with Bayesian PP >0.1068 bfGWAS are colored. (b) Variants with Fgwas PP >0.1068 are colored.

Fig S10. ZoomLocus plots around *rs147859257* in the locus *C3* using the association criteria by P-value, standard BVSR, Fgwas, and bfGWAS. (a) –log10(P-values) by single variant tests. (b) Bayesian PPs by BVSR. (c) Fgwas PPs. (d) Bayesian PPs by bfGWAS. The purple triangle in (b, d) denotes variant *rs147859257*; the blue triangle in (a, c) denotes the top significant variant (*rs2230199*) by P-value.

Fig S11. Fgwas enrichment estimates with 95% error bars, using various functional annotations. (a) Gene-based annotations. (b) Regulatory annotations. (c) Chromatin states profiled in the K562 cell line.

Fig S12. Ratios of enrich parameters versus the respective genome-wide averages, along with 95% confidence intervals, using various functional annotations. (a, c, e) Causal probability ratios *π*_*q*_/*π*_*avg*_. (b, d, f) Effect-size variance ratios 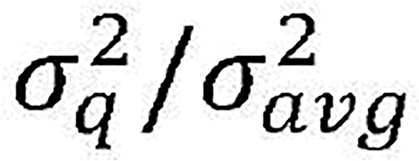.

Fig S13. Enrichment parameter estimates with 95% error bars by bfGWAS and Fgwas, with chromatin states profiled in different cell lines. (a, b, c) H1-hESC cell line. (d, e, f) GM12878 cell line. (g, h, i) HepG2 cell line. Missing bfGWAS estimates are due to none enrichment (estimates are the same as their priors), while missing Fgwas estimates are due to convergence issues.

Fig S14. Enrichment parameter estimates with 95% error bars by bfGWAS and Fgwas, with chromatin states profiled in different cell lines. (a, b, c) HUVEC cell line. (d, e, f) HSMM cell line. (d, h, i) NHLF cell line.

Fig S15. Enrichment parameter estimates with 95% error bars by bfGWAS and Fgwas, with chromatin states profiled in different cell lines. (a, b, c) NHEK cell line. (d, e, f) NHEK cell line.

